# Adapting machine-learning algorithms to design gene circuits

**DOI:** 10.1101/213587

**Authors:** Tom Hiscock

**Affiliations:** Department of Systems Biology, Harvard Medical School, Boston, 02115, USA; Present address: Cancer Research UK, Cambridge Institute, Li Ka Shing Centre, Robinson Way, Cambridge, CB2 0RE, UK.

## Abstract

Biological systems rely on complex networks, such as transcriptional circuits and protein-protein interaction networks, to perform a variety of functions e.g. responding to stimuli, directing cell fate, or patterning an embryo. Mathematical models are often used to ask: given some network, what function does it perform? However, we often want precisely the opposite i.e. given some circuit – either observed *in vivo*, or desired for some engineering objective – what biological networks could execute this function? Here, we adapt optimization algorithms from machine learning to rapidly screen and design gene circuits capable of performing arbitrary functions. We demonstrate the power of this approach by designing circuits (1) that recapitulate important *in vivo* phenomena, such as oscillators, and (2) to perform complex tasks for synthetic biology, such as counting noisy biological events. Our method can be readily applied to biological networks of any type and size, and is provided as an open-source and easy-to-use python module, GeneNet.

## Introduction

Biological networks – sets of carefully regulated and interacting components – are essential for the proper functioning of biological systems (Barabasi and Oltvai, 2004; Zhu et al., 2007). Networks coordinate many different processes within a cell, facilitating a vast array of complex cell behaviors that are robust to noise yet highly sensitive to environmental cues. For example, transcription factor networks program the differentiation of cells into different cell types (Cahan et al., 2014; Goode et al., 2016; Plath and Lowry, 2011), orchestrate the patterning of intricate structures during development (Davidson, 2010; Rhee et al., 2014), and allow cells to respond to dynamic and combinatorial inputs from their external environment (Lopez-Maury et al., 2008; Mangan and Alon, 2003). In addition to transcriptional regulation, many other processes form biological networks, including protein-protein interactions (Stelzl et al., 2005), post-translational modifications (Minguez et al., 2012), phosphorylation (Linding et al., 2007) and metabolism (Fiehn, 2001; Jeong et al., 2000).

Understanding how these networks execute biological functions is central to many areas of modern biology, including cell biology, development and physiology. Whilst the network components differ between these disciplines, the principles of network function are often remarkably similar. This manifests itself as the recurrence of common network designs (“network motifs”) in transcriptional, metabolic, neuronal and even social networks (Alon, 2003, 2007; Milo et al., 2002; Rosenfeld et al., 2002; Shen-Orr et al., 2002). For example, negative feedback is a network design that achieves homeostasis and noise resilience, whether that be in the regulation of glucose levels, body temperature (Simon et al., 1986), stem cell number (Shraiman, 2005), or gene expression levels (Lestas et al., 2010).

A major challenge to understand and ultimately engineer biological networks is that they are complex dynamical systems, and therefore difficult to predict and highly non-intuitive (Alon, 2006). Consequently, for anything other the simplest networks, verbal descriptions are insufficient, and we rely on computational models, combined with quantitative data to make progress. For example, after decades of genetics, biochemistry, quantitative microscopy and mathematical modeling, a fairly complete, and predictive, description of *Drosophila* anterior-posterior patterning is emerging (Fowlkes et al., 2008; Gregor et al., 2007; Jaeger et al., 2004a; Jaeger et al., 2004b; Manu et al., 2009a, b).

Quantitative approaches have also proven useful in the rational design of circuits for synthetic biology (Khalil and Collins, 2010; Mukherji and van Oudenaarden, 2009). If we are to successfully engineer biological processes (e.g. for the production of biomaterials, for use as biosensors in synthetic biology, or for regenerative medicine (Davies, 2017; Khalil and Collins, 2010)), then we need clear design principles to construct networks. This requires formal rules that determine which components to use and how they should interact. Mathematical models, combined with an expanding molecular biology toolkit, have enabled the construction of gene circuits that oscillate (Elowitz and Leibler, 2000), have stable memory (Gardner et al., 2000), or form intricate spatial patterns (Liu et al., 2011).

One approach to analyze and design gene circuits is to propose a network and, through computational analysis, ask whether it (i) fits some observed data, or (ii) performs some desired function; and if not, modify parameters until it can. For example, Elowitz and Leibler (2000) proposed the repressilator circuit, used simulations to show it should oscillate, and demonstrated its successful operation in *E. coli*. This approach critically relies on starting with a “good” network i.e. one that is likely to succeed. How do you choose a “good” network? In the study of natural networks, this can be guided by what is known mechanistically about the system (e.g. from ChIP-seq data). However, often the complete network is either unknown or too complicated to model and therefore researchers must make an educated guess for which parts of the network are relevant. For synthetic circuits, one can emulate natural designs and/or use intuition and mathematical modeling to guide network choice. In both cases, these approaches start from a single network – either based on some understanding of the mechanism, or on some intuition of the researcher, or both – and then ask what function this network performs. (Note, throughout we use the term “function” to refer to two things: (1) some real biological function e.g. patterning the *Drosophila* embryo, or (2) some engineered function in synthetic biology e.g. an oscillator circuit.)

Here we aim to do the exact opposite, namely to ask: given a prescribed function, what network(s) can perform this function? Equivalently, this means considering the most general network architecture possible (i.e. all genes can activate/repress all other genes), and then determining for what parameters (i.e. what strengths of activation/repression) the network executes the desired function. Such numerical screens have discovered a wide range of interesting gene circuits, including: fold change detectors (Adler et al., 2017), robust oscillators (Li et al., 2017), stripe-forming motifs (Cotterell and Sharpe, 2010), polarization generators (Chau et al., 2012), robust morphogen patterning (Eldar et al., 2002), networks that can adapt (Ma et al., 2009), gradients that scale (Ben-Zvi et al., 2008) and biochemical timers (Gerardin and Lim, 2017).

These studies demonstrate that unbiased and comprehensive *in silico* screens of gene circuits can generate novel and useful insights into circuit function. However, the drawback of such an approach is that it is computationally expensive, and becomes prohibitively slow as the network size is increased, due to the high dimensional parameter spaces involved. For example, consider a gene circuit consisting of *N* genes, where each gene can activate or repress any other gene. There are then *N*^2^ interactions in this network i.e. at least *N*^2^ parameters. It is therefore challenging to scan through this high dimensional parameter space to find parameter regimes where the network performs well. There are more efficient algorithms to search through parameter space, many of which randomly change parameters and then enrich for changes that improve network performance. Such approaches include Monte Carlo methods and their extensions (Crombach et al., 2012; Perkins et al., 2006), as well as algorithms that mimic natural selection *in silico* (Francois and Hakim, 2004; Francois et al., 2007). However, these approaches are still computationally intensive and difficult to scale to more complex functions and networks.

Here, we designed an approach inspired by machine learning to significantly accelerate the computational screening of gene circuits. In machine learning, complex models (typically ‘neural networks’) with a large number of parameters (typically millions) are fit to data to perform some prescribed function (Hinton and Salakhutdinov, 2006; LeCun et al., 2015). For example, in computer vision, this function could be to detect a human face in a complex natural scene (Li, 2015). Many of the successes in machine learning have been underpinned by advances in the algorithms to fit parameters to data in high dimensions. Central to these algorithms is the principle of “gradient descent”, where instead of exhaustively screening parameter space, or randomly moving within it, parameters are changed in the direction that most improves the model performance (Amari, 1993). An analogy for gradient descent is to imagine you are walking on a mountain range in the fog and wish to descend quickly. An effective strategy is to walk in the direction of steepest downhill, continuously changing direction as the terrain varies, until you reach the base. Analogously, gradient descent works by iteratively changing parameters in the “steepest” direction with respect to improving model performance.

A major challenge is to efficiently compute these directions in high dimensions. This relies on being able to differentiate the outputs of a complex model with respect to its many parameters. A key advance in this regard has been to perform differentiation *automatically* using software packages such as Theano (Bergstra et al., 2010) and Tensorflow (Abadi et al., 2016). Here, gradients are not calculated using pen and paper, but instead algorithmically, and therefore can be computed for models of arbitrary complexity.

We realized that training neural networks is in many ways similar to designing biological circuits. Specifically, we start with some prescribed function (or data), and we then must fit a model with a large number of parameters to perform the function (fit the data). We thus reasoned that we could use exactly the same tools as in machine learning to design gene circuits, namely advanced gradient-descent, Adam (Kingma and Ba, 2014), to fit parameters, and automatic differentiation with Theano/Tensorflow to calculate gradients. We found that such an approach could effectively and rapidly generate circuits that perform a range of different functions, using a fairly simple python module, “GeneNet”, which we make freely available.

## Results

### Algorithm overview

We consider a generic model of a transcriptional gene circuit that has been proposed and used elsewhere (Cotterell and Sharpe, 2010; Jaeger et al., 2004a; Molinelli et al., 2013). The model comprises *N* transcription factors, whose concentrations are represented by the *N*-component vector, ***y***. We assume that all interactions between genes are possible; this is parameterized by a *N × N* matrix, **W**. Thus each element *W*_*ij*_ specifies how the transcription of gene *i* is affected by gene *j –* if *W*_*ij*_ is positive, then *j* activates *i;* if *W*_*ij*_ is negative, then *j* inhibits *i*. We further assume that each gene is degraded with rate *k*_*i*_. Together this specifies an ordinary differential equation (ODE) model of the networ:

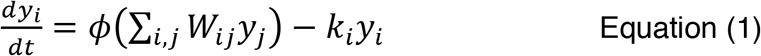

Here, *ϕ*(*x*) is a nonlinear function, ensuring that transcription rates are always positive, and saturate at high levels of the input. The task of network design is thus to find the parameters **W** and **k** such that the network operates as desired.

Note that Equation 1 represents just one specific form of gene circuit model that we focus on here. We emphasize, however, that this particular form is not required for our algorithm to work: any ODE-based model can be fit with the same algorithm. Therefore, one could also consider more realistic models including, for example, epistasis and non-additive effects.

To fit Equation 1, we start with an analogy to neural networks. Neural networks are highly flexible models with large numbers of parameters that are capable of performing functions of arbitrary complexity (Hornik, 1989), whose parameters must be fit to ensure the network performs some designed function. We wondered whether the algorithms used to fit neural network parameters could be adapted to fit gene circuit parameters.

To do this, we start with the simplest example where we wish the network to compute some input-output function, *y* = *f*(*x*). In this case, we allow one of the genes *y*_1_ ≡ *x*, to respond to external input, and examine the output of another gene, *y*_*N*_ ≡ *y*. We then define a “cost”, *C*, which tracks how closely the actual output of the network, *y*, matches the desired output of the network, 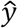. First, this involves specifying what the desired output is; as our first example, we consider the case where we want the network output to respond in an ultrasensitive, switch-like manner to the level of some input, *x*, i.e. *y* = 0 for *x* < *x*_*_ and *y* = 1 for *x* > *x*_*_, as in Figure 1A. Then, we choose the mean squared error as the form of our cost, i.e. 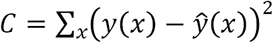.

**Figure 1:**
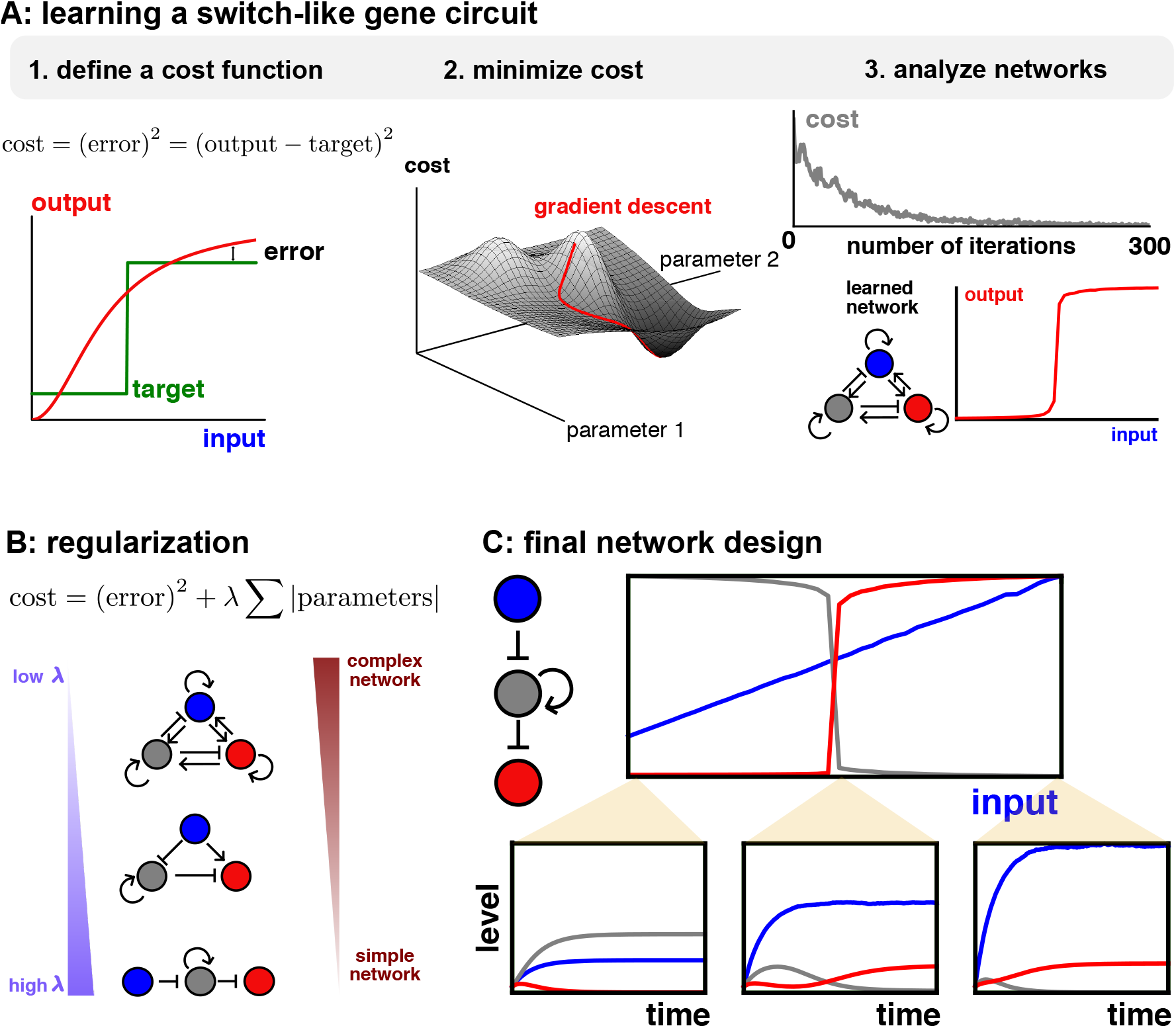
Overview of GeneNet. A: The optimization algorithm consists of three parts: defining a cost function (left), updating parameters to minimize the cost via gradient descent (middle), and analyzing the learned networks (right). B: Regularization selects networks with varying degrees of complexity. C: Final design of an ultrasensitive switch. Upper: the final values of each of the three genes as a function of different levels of input. Lower: time traces illustrating network dynamics for three representative values for the input level.

The goal is then to find the parameters that minimize this cost and therefore specify the network that best gives the desired output. To do this rapidly and efficiently in high dimensional parameter spaces, we use gradient descent. In gradient descent, parameters, *p*_*i*_ are updated in the direction that maximally reduces the cost, i.e. 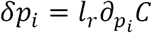, where *l*_*r*_ is the learning rate. Intuitively, for a two-dimensional parameter set, this corresponds to moving directly “downhill” on the cost surface (Figure 1A).

For classic gradient descent, the learning rate *l*_*r*_ is set to a constant value. However, this can present problems when optimizing a highly complex cost function in high dimensions. Intuitively, and as shown in Figure S1, we would like the learning rate to adapt as optimization proceeds (and also to be different for each parameter). A more sophisticated version of gradient descent, Adaptive moment estimation, or Adam (Kingma and Ba, 2014), has been established to overcome these difficulties and is being widely used to fit complex neural network models (Ruder, 2016). For this reason, we choose Adam as our core optimization algorithm; a choice we will later justify.

Minimizing the cost is, in principle, exactly the same whether training neural networks or screening gene circuits. The difference arises when computing the gradient of the cost function with respect to the parameters. For the algebraic equations in neural networks, the computation is fairly straightforward and can be written down explicitly. However, for a gene circuit ODE model, it is highly nontrivial to differentiate the model outputs with respect to circuit parameters. It is, in principle, possible to do by hand, although this is cumbersome and must be repeated anew for each model (Frohlich et al., 2017). An alternative approach is to estimate the gradients empirically, by comparing the outputs of Equation 1 when each of the system parameters is changed by a small amount (a similar approach was used in (Uzkudun et al., 2015)). Again, however, this method scales poorly as the number of parameters to be fit increases.

The key insight is that machine learning libraries, such as Theano and Tensorflow, permit automatic (or “algorithmic”) differentiation of computer programs. Therefore, we implemented a simple differential equation solver in Theano, and using a single line of code, can “differentiate” the solver and thereby compute gradients. (We also tried Tensorflow, but found it to be significantly slower, see Table S3). Together, the gene network model, the cost function, and the gradient descent algorithm define a procedure to design gene circuits (see Methods).

We first tested our pipeline by asking it to generate an ultrasensitive switch, a circuit that is seen *in vivo* (Tyson et al., 2003), and has also been rationally engineered (Palani and Sarkar, 2011). Indeed, we find that as we step through repeated iterations of the gradient descent, we efficiently minimize the cost function, and so generate parameters of a gene network 253 model that responds sensitively to its input (Figure 1A).

To train this circuit, we have used a sophisticated version of gradient descent, Adam. Could a simpler algorithm – namely classic gradient descent with constant learning rate – also work? As shown in Figure S1, we find that we can generate ultrasensitive switches using classic gradient descent, albeit more slowly and after some fine tuning of the learning rate parameter. We emphasize that, in contrast, Adam works well with default parameters. Furthermore, we find that as we consider functions of increasing complexity, classic gradient descent fails to learn, whilst Adam still can. These results echo outputs from the machine learning community, where Adam significantly outperforms other optimization algorithms (Kingma and Ba, 2014).

Whilst the network in Figure 1A performs well, its main drawback is that it is complicated and has many parameters. Firstly, this makes it difficult to interpret exactly what the network is doing, since it is not obvious which interactions are critical for forming the switch. Secondly, it would make engineering such a network more complicated. Therefore, we modified the cost in an attempt to simplify gene networks, inspired by the techniques of “regularization” to simplify neural networks (Goodfellow et al., 2016). Specifically, we find that if we add the L1 norm of the parameter sets to the cost, i.e. 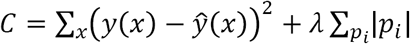, we can simplify networks without significantly compromising their performance. Intuitively, the extra term penalizes models that have many non-zero parameters i.e. more complex models (Brunton et al., 2016). By varying the strength of the regularization, *λ*, we find networks of varying degrees of complexity (Figure 1B).

The final output of our algorithm is a simplified gene network that defines a dynamical system whose response is a switch-like function of its inputs (Figure 1C). Therefore, we have demonstrated that machine-learning algorithms can successfully train gene networks to perform a certain task. In the remainder of this work, we show the utility of our pipeline by designing more realistic and complex biological circuits.

## Applications

First, we consider three design objectives for which there already exist known networks, so that we can be sure our algorithm is working well. We find that we can rapidly and efficiently design gene circuits for each of the three objectives by modifying just a few lines of code that specify the objective, and without changing any details or parameters of the learning algorithm. Further, we can screen for functional circuits within several minutes of compute time on a laptop. In each case, the learned network is broadly similar to the networks described previously, lending support to our algorithm.

### French-Flag circuit

The first design objective is motivated from the French-Flag model of patterning in developmental biology (Wolpert, 1969). Here, a stripe of gene expression must be positioned at some location within a tissue or embryo, in response to the level of some input. This input is typically a secreted molecule, or “morphogen”, which is produced at one location, and forms a gradient across the tissue. In order to form a stripe, cells must then respond to intermediate levels of the input. To identify gene circuits capable of forming stripes, we ran our algorithm using the desired final state as shown in Figure 2A. The algorithm converges on a fairly simple network, where the input directly represses, and indirectly activates, the output, thus responding at intermediate levels of the input (Figure 2A). Exactly the same network was described in a large-scale screen of stripe-forming motifs (Cotterell and Sharpe, 2010), and has been observed in early *Drosophila* patterning (Clyde et al., 2003), suggesting that our learned design may be a common strategy.

**Figure 2:**
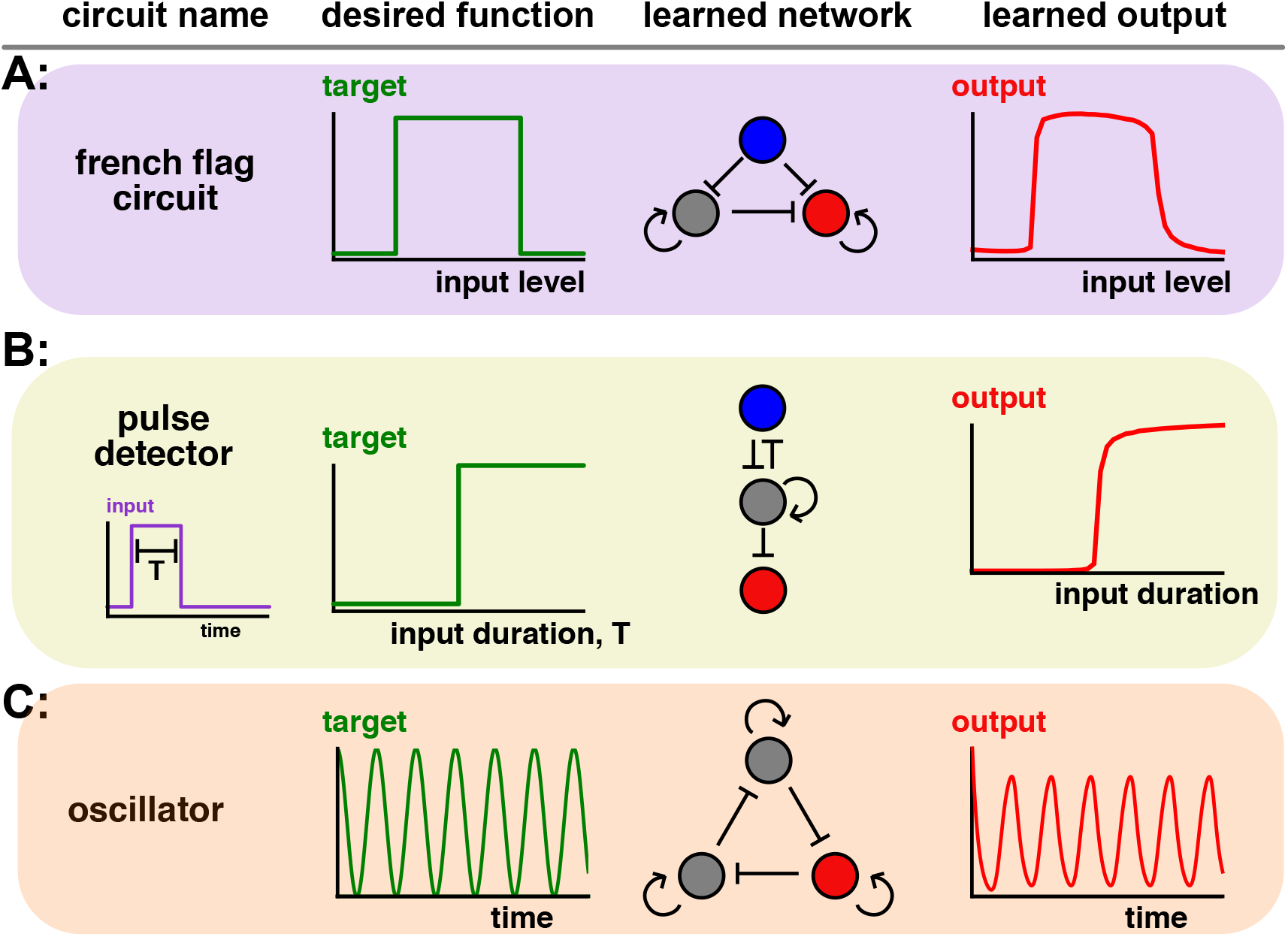
Using GeneNet to learn gene circuits. A: A French-Flag circuit responds to intermediate levels of input to generate a stripe. The red node corresponds to the output gene, and the blue node the input. B: A “duration-detector” which is irreversibly activated when stimulated by pulses exceeding a certain critical duration. As above, the red node corresponds to the output gene, and the blue node the input. C: An oscillator.

### Pulse detection

In our second example, we consider a more complicated type of input, namely pulses of varying duration. In many cases, cells respond not just to the level of some input, but also to the duration (Hopfield, 1974; Mangan et al., 2003). We sought a circuit design to measure duration, such that once an input exceeding a critical duration is received, the output is irreversibly activated (Figure 2B). As before, by changing a few lines of code, and within a few minutes of laptop compute time, we can efficiently design such a circuit (Figure 2B). This circuit shares features (such as double inhibition and positive feedback) with networks identified in a comprehensive screen of duration detection motifs (Gerardin and Lim, 2017).

### Oscillator

Our third example takes a somewhat different flavor, where instead of training a network to perform a specific input/output function, we train a network to self-generate a certain dynamical pattern – oscillations (Novak and Tyson, 2008; Stricker et al., 2008). In this case, the cost is not just dependent on the final state, but on the entire dynamics, and is implemented by the equation 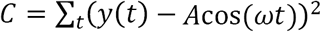, where *A* is the amplitude and *ω* the frequency of the oscillator. Minimizing this cost yields a network that gives sustained oscillations, and is reminiscent of the repressilator network motif that first demonstrated synthetic oscillations (Elowitz and Leibler, 2000). Interestingly, when plotting how the cost changed in the minimization algorithm, we saw a precipitous drop at a certain point (Figure S2), demonstrating visually the transition through a bifurcation to produce oscillations.

### A more complex circuit: a robust biological counter

First, to illustrate the scalability of GeneNet, we considered larger networks (up to 9 nodes, with 81 parameters) and asked whether they could be successfully screened. As shown in Figure S2, we find that the median number of iterations required to train a French Flag circuit is largely insensitive to the network size, demonstrating that GeneNet is scalable to models of increased complexity.

In our final example, we attempt a more ambitious design objective – a biological counter – to demonstrate that GeneNet can also design circuits to perform more complex computations and functions. Whilst counters are found in some biological systems (such as in telomere length regulation (Marcand et al., 1997)), we focus our aim on designing a counter for synthetic biology. There are numerous applications for such counters, two of which are: (1) a safety mechanism programming cell death after a specified number of cell cycles, (2) biosensors that non-invasively count the frequency of certain stimuli, particularly low frequency events (Friedland et al., 2009).

We consider an analog counter, where we wish some input set of pulses to result in an output equal (or proportional) to the number of pulses (Figure 3A). For example, the “input” here, could be the level of a cell cycle related protein to count divisions (Slomovic et al., 2015). As is shown in Figure 3B, the simplest way to count would be simply to integrate over time the levels of the input. One implementation of this used an analog memory device driven by CRISPR mutagenesis (Perli et al., 2016), i.e. when the stimulus is present, Cas9 is active and mutations arise. However, a major shortcoming of such a circuit is that it is unreliable and sensitive to variations in pulse amplitude and duration that are often present (Figure 3B).

**Figure 3:**
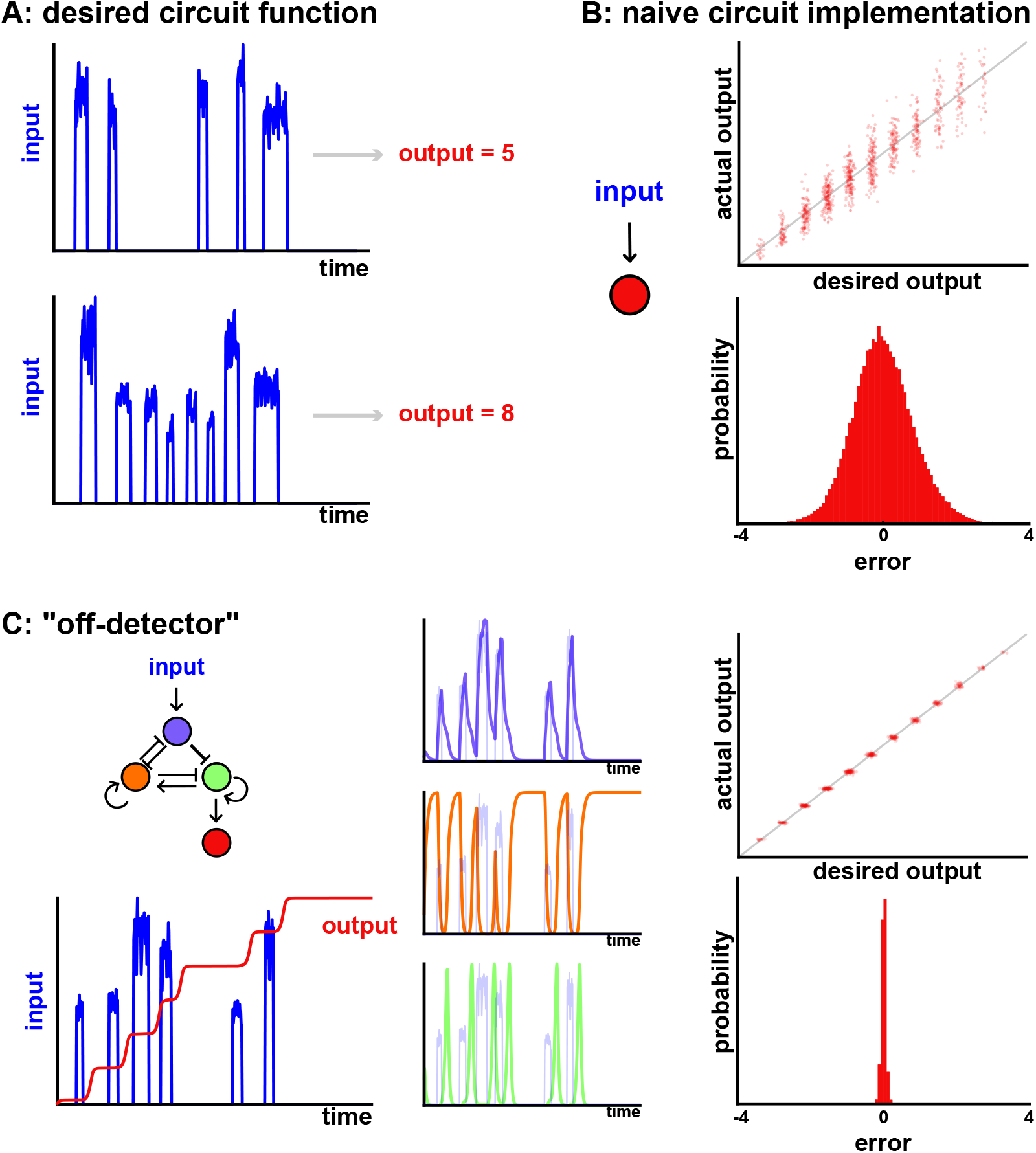
Designing a robust biological counter using GeneNet. A: Desired input/output function of a robust biological counter. B: Simply integrating the input over time yields an analog counter with significant error, as shown by the spread in the input/output function (upper) and the relative errors between the two (lower). C: GeneNet learns an “off-detector” network (left), with substantially reduced error rates (right). Inspection of gene dynamics (middle) shows that an excitable “digital” pulse of green activity is initiated at the end of the input pulse (shaded blue line).

Therefore, we sought to systematically design a novel gene circuit to count pulses that would be robust to their amplitude and duration. To do this, we provided a complex ensemble of input stimuli, each containing a different number of pulses, of varying amplitudes and durations. For each input we then defined the desired output to be equal to the number of pulses present, and trained the network to minimize the mean squared error cost, as before. Strikingly, this procedure uncovers a network that is highly robust in counting pulse number (Figure 3C). (We also report the design of a simpler network that counts pulses, Figure S3, albeit less efficiently that in Figure 3C).

Looking more deeply into the network, we see that it has learned a very interesting way to count, with two key ingredients. Firstly, it acts as an “off-detector”. Specifically, examining the dynamic time traces reveals that the network responds *after* the pulse has occurred, as it turns off. Mechanistically, when the input increases, this allows build up of the purple node and repression of the orange node. However, activation of the downstream green node is only possible once the input pulse has ended, and the repression from the purple node has been alleviated. In this way, the circuit responds to the termination of the pulse, and is thus robust to its duration.

Secondly, the network uses “digital encoding” to be robust to the level of the input. This is achieved by having the green node undergo an “excitable pulse” of stereotyped amplitude and duration, which is then integrated over time by the red node to complete the counter. By using “digital” pulses of activity with an excitable system, the circuit is therefore insensitive to the precise levels of the input. Together, this forms a circuit that reliably counts despite large variations in input stimulus.

We emphasize that these behaviors have not been hard-coded by rational design, but rather have emerged when training the network to perform a complex task. This example therefore shows that a more challenging design objective can be straightforwardly accommodated into our gene network framework, and that it is possible to learn rather unexpected and complex designs.

## Discussion

In this work, we have developed an algorithm to learn gene circuits that perform complex tasks (e.g. count pulses), compute arbitrary functions (e.g. detect pulses of a certain duration) or resemble some real biological phenomenon (e.g. a French-flag circuit). We have demonstrated that these networks can be trained efficiently on a personal laptop, and require neither fine-tuning of algorithm parameters nor extensive coding to adapt to different network designs. This ease-of-use means that researchers addressing questions in basic biology can quickly generate models and hypotheses to explain their data, without investing a lot of time carefully building and simulating specific models. Equally, our approach should also allow synthetic biologists to rapidly generate circuit designs for a variety of purposes.

Here we have focused on gene networks and transcriptional regulation as a proof of principle. However, our algorithm is rather general and could easily be extended to learn other networks, such as phosphorylation, protein-protein interaction and metabolic networks, so long as they are described by ordinary differential equations. Further, whilst the networks we have focused on are relatively small and have been trained on a personal laptop, our Theano pipeline can be easily adapted to run much faster on GPUs, and therefore we expect that large networks could also be trained effectively (Bergstra et al., 2010).

Our approach could also be extended to incorporate other types of differential equation, such as partial differential equations. This would allow us to understand how networks operate in a spatial, multicellular context, throughout an entire tissue, and thus provide useful insights into how different structures and patterns are formed during development (Davies, 2017). Other extensions would be to use real data as inputs to the learning algorithm, in which case more sophisticated algorithms would be required to deal with parameter uncertainty (Calderhead et al., 2009; Liepe et al., 2014)

One drawback of our approach is that it selects only a single gene circuit out of many, and thus may ignore alternative circuits that may also be useful or relevant. A natural extension would therefore be to combine the speed of GeneNet’s parameter optimization with a comprehensive enumeration of different network topologies, thus generating a complete ‘atlas’ of gene circuits (Cotterell and Sharpe, 2010).

Finally, one concern with machine learning methods is that the intuition behind the models is hidden within a “black box” and opaque to researchers i.e. the machine, not the researcher, learns. We would like to offer a slightly different perspective. Instead of replacing the researcher, our algorithm acts as a highly efficient way to screen models i.e. one shouldn’t view it as a tool to solve problems, but rather as an efficient way to generate new hypotheses. The role of the scientist is then to: (1) cleverly design the screen (i.e. the cost) such that the algorithm can effectively learn the desired function, and (2) to carefully analyze the learned circuits and the extent to which they recapitulate natural phenomena. The distinct advantage of our approach over neural networks is that we learn real biological models – gene circuits – which are both directly interpretable as mechanism, and provide specific assembly instructions for synthetic circuits.

## Methods

### Gene network model

We considered the following set of coupled ODEs as our gene circuit model, motivated by (Cotterell and Sharpe, 2010; Jaeger et al., 2004a; Jaeger et al., 2004b):

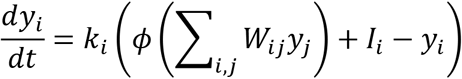

Here, *y*_*i*_ denotes the concentration of gene *i*, where *i* = 1 … *N* for an N-node network. *W*_*ij*_ is a matrix correspond to network weights: *W*_*ij*_ > 0 means that gene *i* activates gene *j*, and *W*_*ij*_ < 0 means that gene *i* represses gene *j*. *k*_*i*_ is a vector that represents the (assumed linear) degradation of gene *i. l*_*i*_ is the prescribed input to the system, which we assume directly causes transcription of gene *i*. The function *ϕ* is a nonlinearity chosen such that the gene network saturates; we choose *ϕ*(*x*) = 1/(exp(*x*) - 1), as in (Cotterell and Sharpe, 2010). 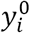 represent the initial conditions of the system. Note, this is a non-dimensionalized version of Equation 1, whereby gene concentrations are normalized to their maximal level.

### Algorithm details

We coded the algorithm using python v2.7.5, and the machine-learning library, Theano v0.8.2, which performs static optimizations for speed and permits automatic differentiation.

The key steps in the algorithm are:

#### 1. Define an ODE Solver

We choose a simple Euler integration method, whereby the differential equation 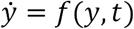 is solved iteratively: 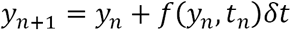. We set *δt* = 0.01. We implement the solver using the theano.scan feature, which optimizes the computation of loops.

#### 2. Define the desired function of the network

This consists of two items: (1) a collection of different inputs to train the network on, and (2) for each input, a desired output. For example, in Figure 2B, we must include inputs of varying durations, and a function that computes whether the pulse exceeds some critical duration. The input can be static in time (as in Figure 2A), dynamic in time (as in Figure 2B) or zero (as in Figure 2C). The desired output can be the final state of the network (as in Figure 2B), or the complete time dynamics of the network (Figure 2C). See Table S1 for further details.

We first tested our pipeline by asking it to generate an ultrasensitive switch, a circuit that is seen *in vivo* (Tyson et al., 2003), and has also been rationally engineered (Palani and Sarkar, 2011). Indeed, we find that as we step through repeated iterations of the gradient descent, we efficiently minimize the cost function, and so generate parameters of a gene network model that responds sensitively to its input (Figure 1A).

#### 3. Define the cost to be minimized

As discussed in the main text, we use the mean squared error as the cost. Since we are often not concerned with absolute levels of any gene, but rather the relative levels, we modify the cost such that the network output can be rescaled by a factor *A*, which is a learnable parameter, i.e. *y → Ay.* For regularization, we add a term 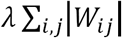 to the *cost*, with the aim of simplifying the network matrix *W*_*ij*_.

#### 4. Define the parameters that are to be fit

For all simulations, we fit the network weights, *W*_*ij*_ to the data. For Figures 1, 2A, 2B, we do not allow *k*_*i*_ to change, and instead set *k*_*i*_ = 1. For Figure 2C and 3C, we use the *k*_*i*_ as learnable parameters. For Figure 2C, we must allow the initial conditions of the network to be learned, such that the oscillator has the correct phase.

#### 5. Define the optimization algorithm

We use the Adam optimizer (Kingma and Ba, 2014) with parameters *l*_*r*_ = 0.1, *b*_1_ = 0.02, *b*_2_ = 0.001, *e* = 10^−8^ in all cases.

#### 6. Initialize the model parameters

We set *k*_*i*_ to be one and the network weights to be a normally distributed random number with mean zero and standard deviation 0.1. Initial conditions (except for Figure 2C) are set as 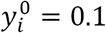.

#### 7. Train the network

We iteratively perform the optimization procedure to update parameters, initially setting *λ* = 0. At each step, we train the network using a subset of total input/output data, using a “batch” of *B* input/output pairs. The idea is that batching the data in this way adds stochasticity to the optimization and therefore avoids local minima.

#### 8. Regularize the network

Once a network has been trained, we now regularize it by increasing *λ* and reimplementing the optimization procedure. A range of different *λ* values is used until a network of desired complexity is achieved.

#### 9. “Prune” the network

In the final step, we retrain a simplified network. Specifically, starting from the regularized network, we set any small network weights to be exactly zero, i.e. 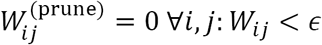, and then optimize over the remaining weights, using *λ* = 0.

#### 10. Save parameters

See Table S1 for the parameter values for the figures used in the main text.

### Implementation details

All computation was performed on a Macbook air, 1.3GHz Intel Core i5, with 8GB 1600 MHz DDR3 RAM. We repeated the learning algorithm for each of the designs in the paper several times, with different regularization levels *λ*, and found similar network topologies to be learned in each case. In the figures we report a representative network, where *λ* has been chosen manually to give a minimal network that still performs the function well. Table S2 gives details of the algorithm implementation specific to each figure.

### Software

Theano is available for download here http://deeplearning.net/software/theano/. GeneNet is available on github: “https://github.com/twhiscock/GeneNet-“.

## Acknowledgements

I thank John Ingraham for inspiring this project over breakfast and dinnertime conversations, and for his enthusiasm and generosity in teaching me machine learning.

---

**Figure S1:**
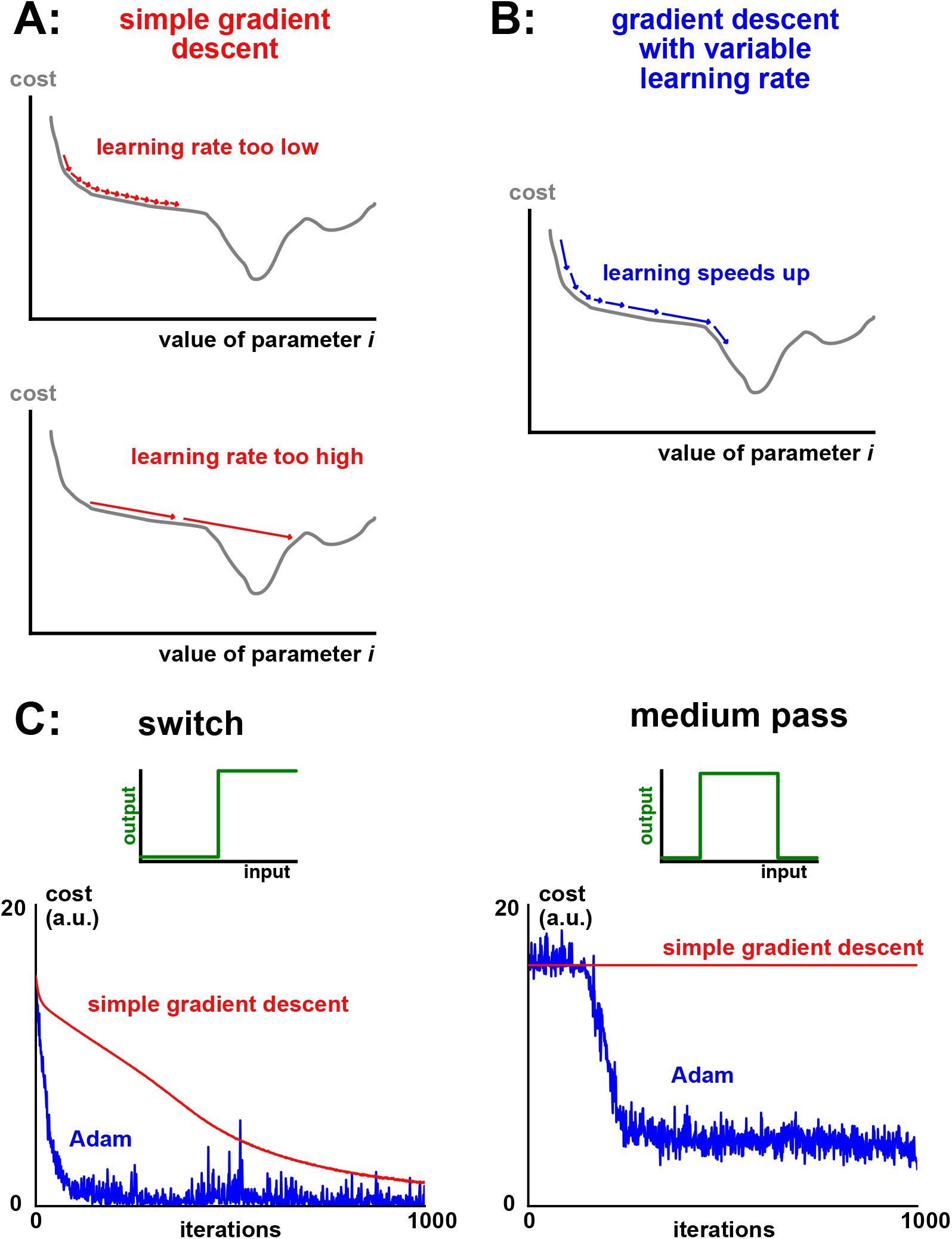
Adam is an effective gradient descent algorithm for ODEs. A: Using a constant learning rate in gradient descent creates difficulties in the optimization process. If the learning rate is too low (upper schematic), the algorithm ‘gets stuck’ in plateau regions with a shallow gradient (saddle points in high dimensions). If instead the learning rate is too high (lower schematic), important features are missed and/or the learning algorithm won’t converge. B: An adaptive learning rate substantially improves optimization (schematic). Intuitively, learning speeds up when traversing a shallow, but consistent, gradient. C: Cost minimization plotted against iteration number, comparing classic gradient descent (red) with the Adam algorithm (blue). Left: an ultrasensitive switch. Right: a medium pass filter.

**Figure S2:**
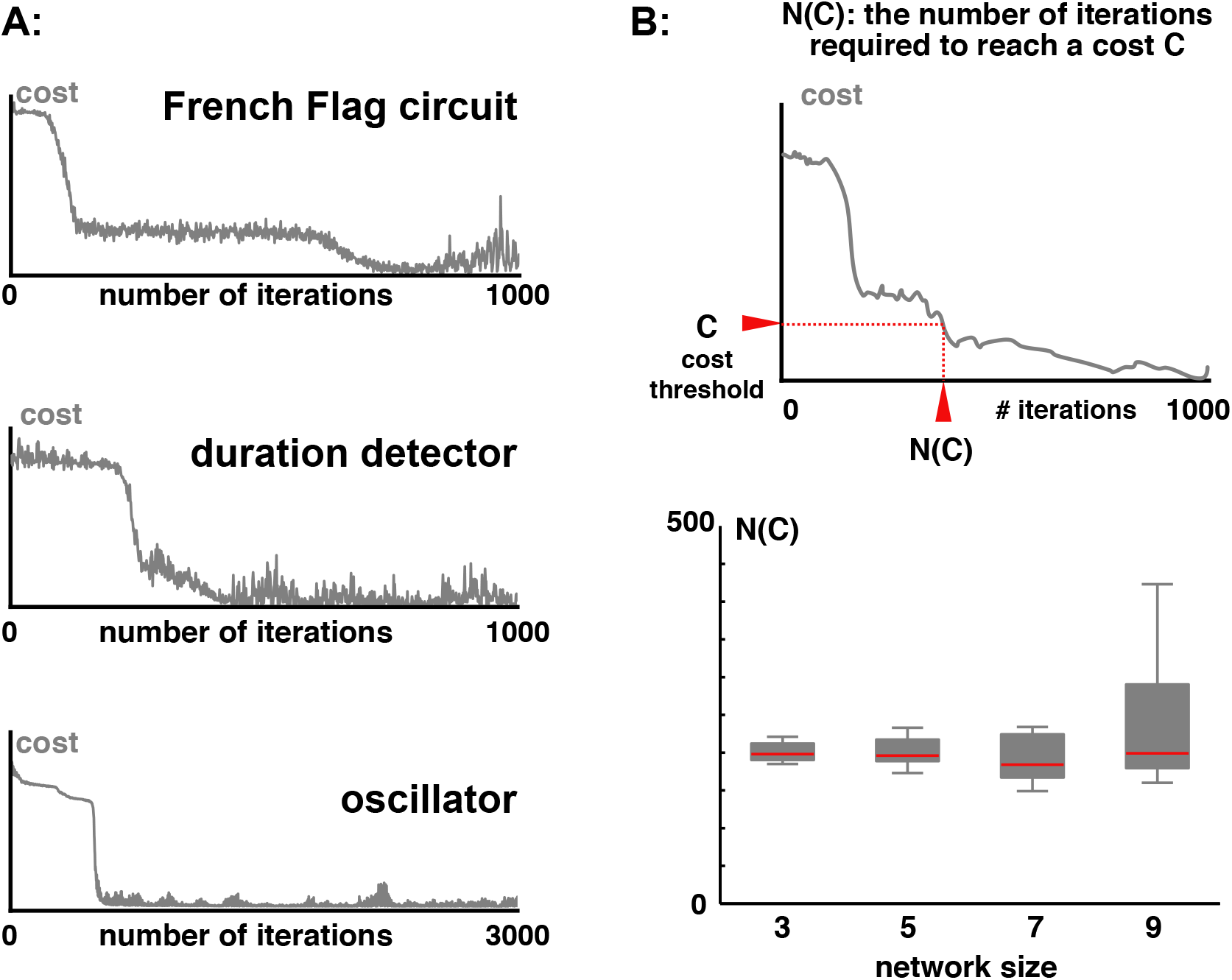
Cost minimization. A: Example traces of the cost minimization during the optimization procedure. Note, in all cases (and particularly in the oscillator), there are sharp drop-offs in cost, which are likely reflecting bifurcation points in the dynamics. B: Scalability of GeneNet. Upper: schematic of N(C) – the number of iterations required to reach a given network performance. Lower: For the same desired circuit function as in Figure 2A, we train networks of varying sizes and provide a boxplot of the number of iterations required to achieve a cost, *C = 0.4*. The median is roughly constant.

**Figure S3:**
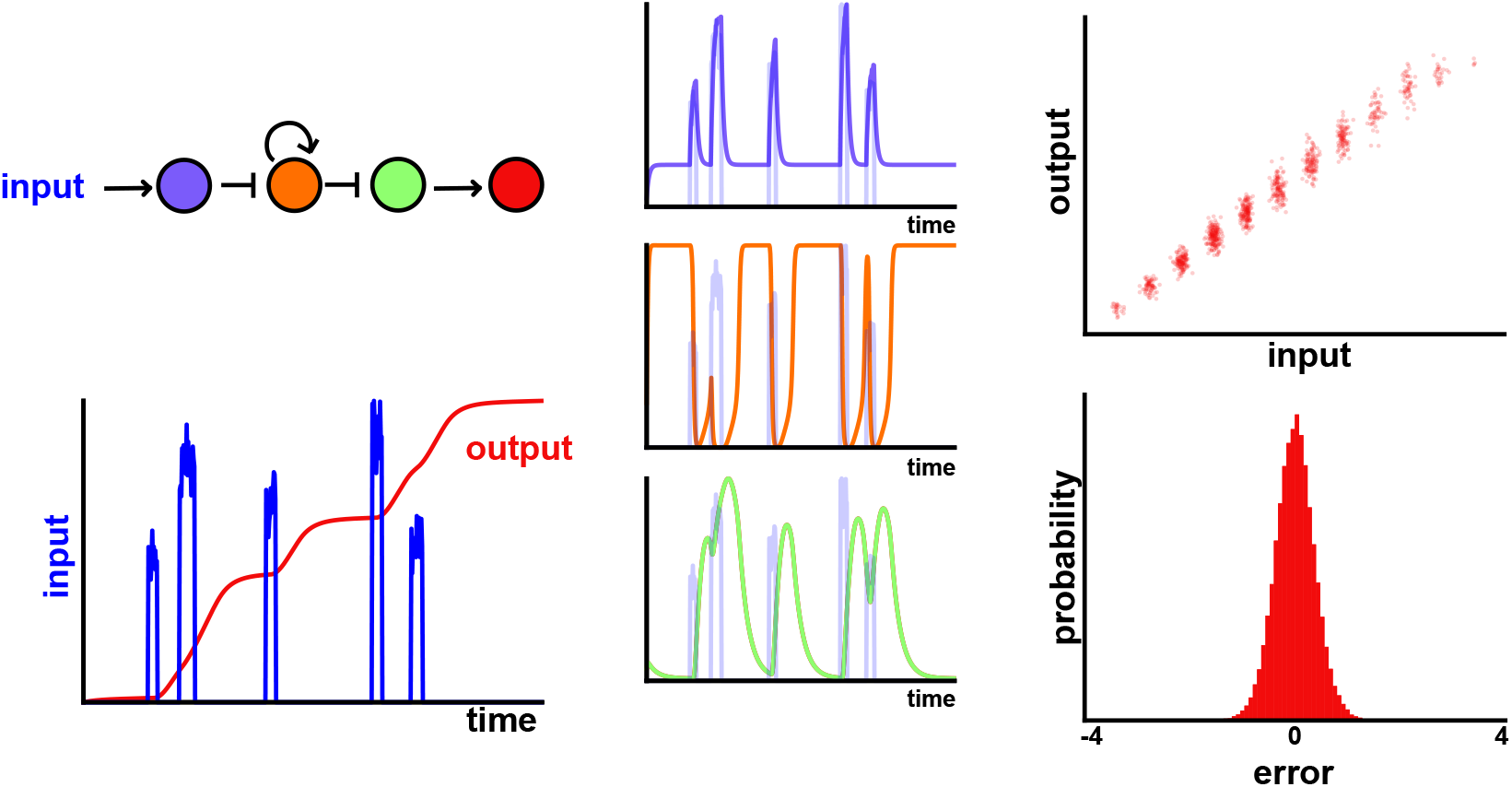
A simpler counter circuit. By using a larger regularization parameter, *λ*, than in the main text, we generate a simpler, but less reliable counter circuit.

**Table S1:** Parameter values for networks learned in the main text.

| Network | Input | Desired output | Learned $W_{ij}$ | Learned $k_i$ |
| --- | --- | --- | --- | --- |
| **Switch**<br>**(Fig 1)** | $x(t) = A$<br>$A \sim U(0,2)$<br>(U is the uniform random distribution) | $\hat{y} = 1$ for $x > 1$<br>$\hat{y} = 0$ for $x < 1$ | $\begin{pmatrix} 0 & 0 & 0 \\ -5.5 & 12.9 & 0 \\ 0 & -5.6 & 0 \end{pmatrix}$ | $\begin{pmatrix} 1 \\ 1 \\ 1 \end{pmatrix}^*$ |
| **French-flag**<br>**(Fig 2A)** | $x(t) = A$<br>$A \sim U(0,2)$ | $\hat{y} = 1$ for $0.5 < x < 1$<br>$\hat{y} = 0$ otherwise | $\begin{pmatrix} 0 & 0 & 0 \\ -6.3 & 10.8 & 0 \\ -2.2 & -5.8 & 10.0 \end{pmatrix}$ | $\begin{pmatrix} 1 \\ 1 \\ 1 \end{pmatrix}^*$ |
| **Duration**<br>**(Fig 2B)** | $x(t) = 1$ for $t = \left[\frac{1}{6}, \frac{1}{6} + \Delta T\right]$<br>$x(t) = 0$ otherwise<br>$\Delta T \sim U(0, \frac{1}{3})$ | $\hat{y} = 1$ for $\Delta T > \frac{1}{6}$<br>$\hat{y} = 0$ otherwise | $\begin{pmatrix} 0 & -6.1 & 0 \\ -7.8 & 6.1 & 0 \\ 0 & -5.0 & 0 \end{pmatrix}$ | $\begin{pmatrix} 1 \\ 1 \\ 1 \end{pmatrix}^*$ |
| **Oscillator**<br>**(Fig 2C)** | $x(t) = 0$ | $\hat{y}(t) = 1 + \cos(\omega t)$ | $\begin{pmatrix} 3.6 & -3.6 & 0 \\ 0 & 3.2 & -3.7 \\ -6.3 & 0 & 5.1 \end{pmatrix}$ | $\begin{pmatrix} 3.3 \\ 7.3 \\ 1.0 \end{pmatrix}$ |
| **Counter**<br>**(Fig 3)** | $N$ pulses,<br>Amplitude: $A \sim U(1,2)$,<br>Duration: $\Delta T \sim U(10\delta t, 30\delta t)$<br>Separation: $\delta T \sim \exp(1/\lambda)$<br>Pulse rate: $\lambda \sim U(0, 100\delta t)$ | $\int_0^T \hat{y}(t) dt = N$ | $\begin{pmatrix} 0 & -4.0 & 0 \\ -7.2 & 3.2 & 4.1 \\ -2.5 & -10.2 & 6.2 \end{pmatrix}$ | $\begin{pmatrix} 17.6 \\ 27.3 \\ 28.5 \end{pmatrix}$ |

**Table S2:** Algorithm implementation parameters. N: number of iterations B: batch size *λ*: regularization parameter *ϵ*: pruning parameter Note that in some cases we perform multiple optimization procedures, starting with a small batch size to increase noise in the early steps. From our experimentation, the iteration number, N, and batch size, B, can be varied significantly and the algorithm still works – one simply needs to increase *N* and *B* so that enough training samples are used. Here we report the parameters we used to generate the figures in the main text, but expect the precise values to be unimportant, and should be chosen/modified by the user.

| Network | Training | Regularization | Pruning |
| --- | --- | --- | --- |
| **Switch**<br>**(Fig 1)** | $N = 400$ | $N = 1000$ | $N = 1000$ |
| | $B = 128$ | $B = 128$ | $B = 128$ |
| | $\lambda = 0.0$ | $\lambda = 0.56$ | $\lambda = 0.0$ |
| | | | $\epsilon = 1$ |
| **French-flag**<br>**(Fig 2A)** | $N = 1000, B = 2,$ | $N = 1000$ | $N = 1000$ |
| | $N = 1000, B = 128,$ | $B = 2$ | $B = 64$ |
| | $\lambda = 0.0$ | $\lambda = 0.02$ | $\lambda = 0.0$ |
| | | | $\epsilon = 1$ |
| **Duration**<br>**(Fig 2B)** | $N = 1000$ | $N = 1000$ | $N = 1000$ |
| | $B = 64$ | $B = 2$ | $B = 64$ |
| | $\lambda = 0.0$ | $\lambda = 0.02$ | $\lambda = 0.0$ |
| | | | $\epsilon = 1$ |
| **Oscillator**<br>**(Fig 2C)** | $N = 1000$ | $N = 400$ | $N = 1000$ |
| | $B = 1$ | $B = 1$ | $B = 1$ |
| | $\lambda = 0.0$ | $\lambda = 0.02$ | $\lambda = 0.0$ |
| | | | $\epsilon = 1$ |
| **Counter**<br>**(Fig 3)** | $N = 10000, B = 2,$ | $N = 1000$ | $N = 1000$ |
| | $N = 3000, B = 128,$ | $B = 64$ | $B = 64$ |
| | $N = 10000, B = 64,$ | $\lambda = 0.78$ | $\lambda = 0.0$ |
| | $\lambda = 0.0$ | | $\epsilon = 1$ |

**Table S3.** Speed tests. Compute time on Macbook air, 1.3GHz Intel Core i5 CPU, with 8GB 1600 MHz DDR3 RAM for different software implementations. Given is the time to perform 1000 iterations of the learning algorithm, with a batch size of 64, simulating 300 timesteps. We expect speed to be significantly improved using GPUs.

| **Implementation** | **Compute time (s)** |
| --- | --- |
| **Theano**<br>**(unoptimized)** | 48.5 |
| **Theano**<br>**(compiled for optimization)** | 28.5 |
| **Tensorflow** | 309.0 |

